# Microbial Community Structure and Diversity of Shrimp Paste at Different Fermentation Stages

**DOI:** 10.1101/334136

**Authors:** Lingying Dai, Limei Wang, Jiang Sun, Lixue Zheng, Bin Qi

## Abstract

High-throughput sequencing was used to reveal the highly diverse bacterial populations in shrimp paste at different fermentation stages. We studied three stages of fermentation and obtained 448,916 reads. Using this approach, we revealed the presence of 30 phyla, 55 classes, 86 orders, 206 families and 695 genera of bacteria in the shrimp paste. Shrimp paste in fermentation metaphase had a more diverse microbiota than that in fermentation prophase and fermentation anaphase. Diversity appeared greatest in fermentation anaphase. The four dominant phyla were *Proteobacteria, Firmicutes, Actinobacteria*, and *Bacteroidetes*. The most common genera were *Psychrobacter, Halomonas, Bacillus, Alteribacillus, and Lactococcus.* Their content varied at different stages of fermentation. All the microbiome presented a variety of changes in the microbial diversity of shrimp paste.

**Importance:** Most research on the microbial diversity of shrimp paste has focused on the shrimp culture environment, or the chemical composition and sensory attributes of the paste. Little research has been conducted on the microbial diversity and composition of shrimp paste. The relationship between microbes and the flavor and quality of shrimp paste has thus been unknown. We therefore analyzed the microbial composition and variation of shrimp paste at different stages of fermentation. The dominant bacteria in fermentation prophase, metaphase, and anaphase were identified. Our preliminary findings give some insight into which microbes contribute to the flavor of shrimp paste and suggest how to improve its flavor. In addition, our findings are relevant to optimizing the production of shrimp paste and guaranteeing its quality and safety.

## Introduction

Shrimp paste is widely consumed as a condiment and used as an ingredient throughout China and the south-east Asian region (1–2). It is normally produced by fermenting small shrimp (*Acetes vulgaris*) with salt at a ratio of 5:1 (shrimp to salt, w/w). The mixture is thoroughly blended or homogenized before being compacted in a container. Shrimp paste is rich in protein, calcium, carotenoids, and chitin (3–4). It exhibits anti-oxidant activity (5), lowers cholesterol and blood pressure, and enhances the body’s immune response and other biological activity (6). It thus has great potential as a functional food. Shrimp paste produced using different fermentation technologies has different fermentation cycles. Some pastes are fermented for 3-6 months (7), some for 2 months (8) and some for only 1 month, which is traditional in China (9). We took 1 month as a study period, and divided it into three fermentation stages: prophase, metaphase, and anaphase, each of which was 10 days long.

High-throughput sequencing allows the simultaneous sequencing of millions of DNA molecules, providing detailed and comprehensive analysis of the transcriptome and genome of species (10). Compared to traditional sequencing methods, such as the analysis of 16S rRNA genes through denaturing or temperature gradient gel electrophoresis (DGGE/TGGE) (11), single-stranded conformation polymorphisms (SSCP) (12) and Sanger sequencing (13), high-throughput sequencing has been widely used in the analysis of the microbial populations of many fermented foods, including Fen liquor, cheese, and kefir grains (14–17) due to its advantages of high read length, high accuracy, high throughput, and unbiasedness (18).

Naturally fermented seafood usually contains microorganisms related to the raw materials and the growth environment of the seafood (19). Bacteria are the main microorganisms responsible for seafood fermentation. Research on the microbial diversity of shrimp paste has mostly concentrated on the culture environment of shrimp (20–22) and the chemical composition and sensory attributes of shrimp paste (23). There has been little research into the microbial community structure and diversity in shrimp paste. Studying the microbial diversity at different stages of the fermentation of shrimp paste is of great significance for the dissemination of shrimp paste knowledge, the predictability of the flavor of shrimp paste, and the control of shrimp paste production.

## RESULTS

### Sequencing and bioinformatic analysis

DNA was extracted from shrimp paste at 11 stages of fermentation. Following total genomic DNA extraction, amplicons of V3-V4 16S rRNA genes were generated, and 448,916 reads were obtained through high-throughput sequencing, corresponding to 153,766 reads from fermentation prophase, 169,915 reads from fermentation metaphase and 125,235 reads from fermentation anaphase. Species diversity and richness were calculated for each time point (Table 1). ACE values and Chao1 values reflect community richness. Shannon values, Simpson values, and coverage values reflect community diversity. Shrimp paste in fermentation prophase had a less diverse microbiota than that in fermentation metaphase. Diversity appeared greatest in Stage 9 in late fermentation, but it reduced again in the next stage.

**TABLE 1:**
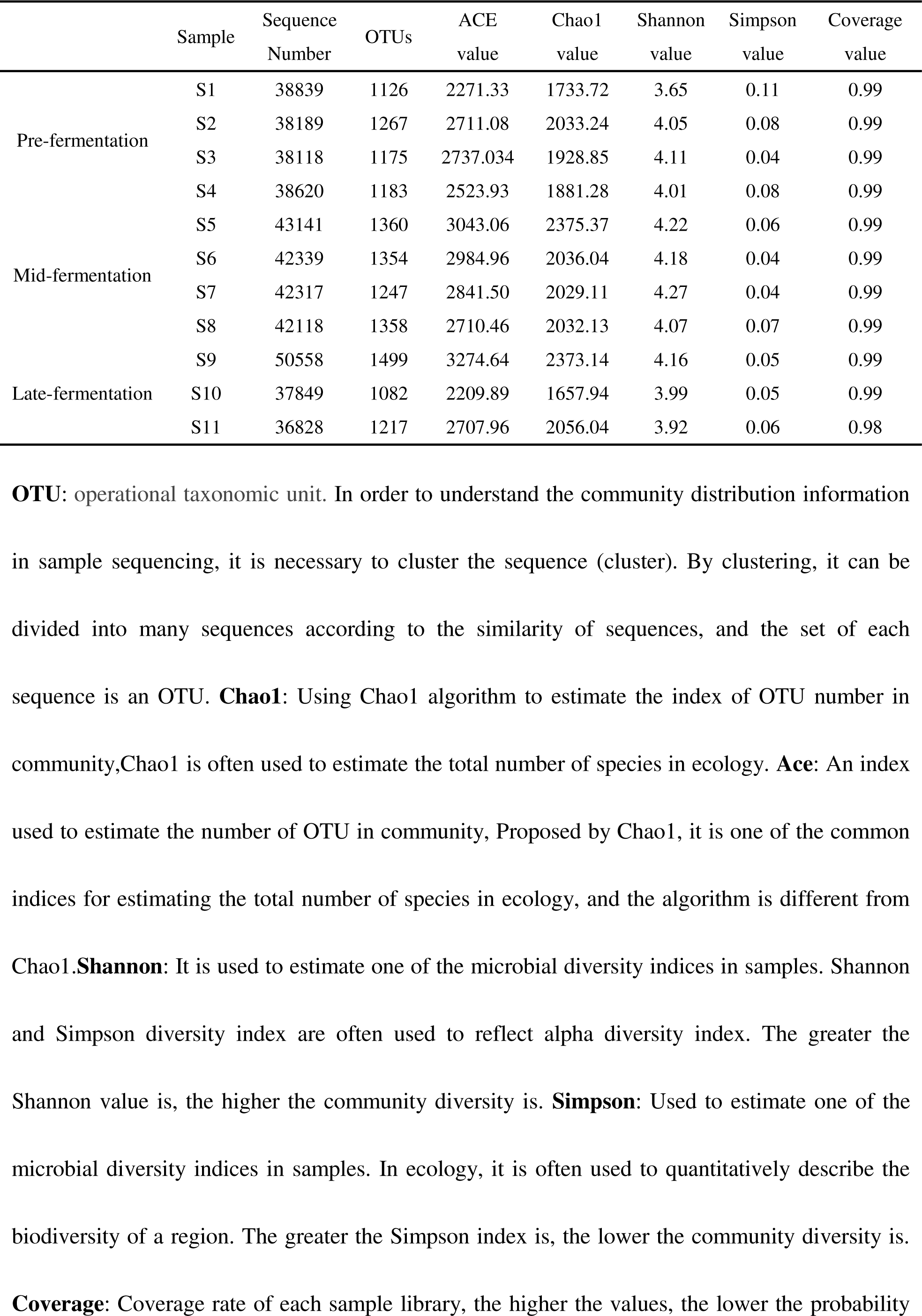
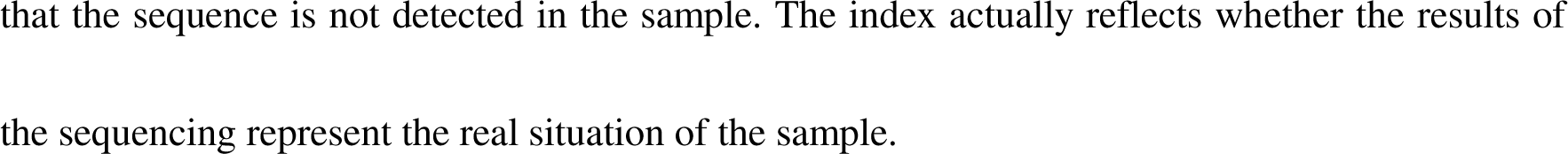
Sequencing richness, diversity and coverage of shrimp paste

### Microbial composition of shrimp paste as revealed by high-throughput sequencing

Phylogenetic assignment (Fig. 1) of high-throughput sequence data revealed the presence of bacteria mainly corresponding to four phyla: *Proteobacteria, Firmicutes, Actinobacteria*, and *Bacteroidetes*. As can be seen in Figure 1, *Proteobacteria* was dominant in the fermenting shrimp paste, reaching 55.59%. The content of *Firmicutes* was slightly lower than that of *Proteobacteria*, accounting for 35.3%. *Actinobacteria* and *Bacteroidetes* were present at the lowest levels (3.09% and 4.89%, respectively).

**Figure 1:**
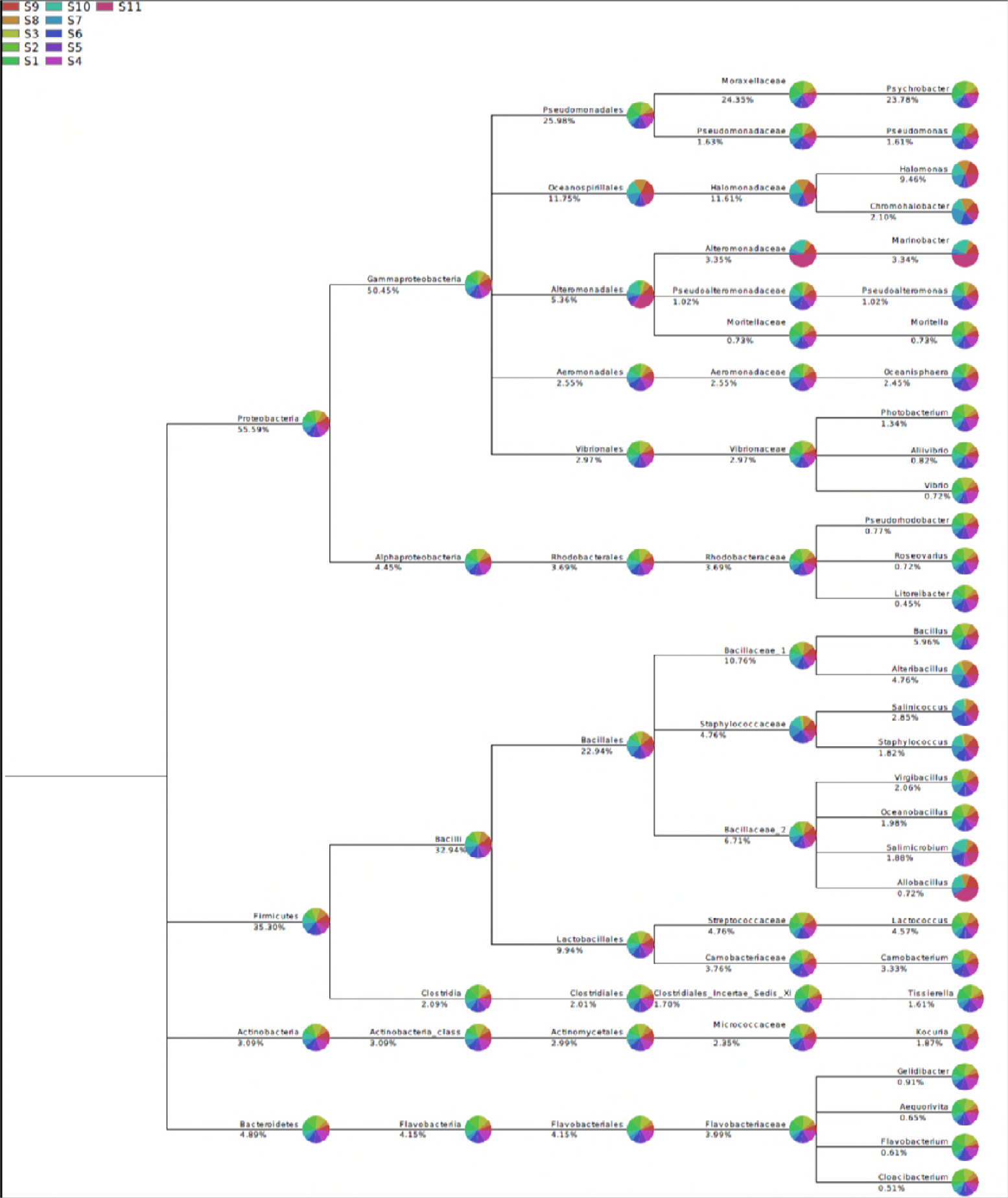
Phylogenetic tree of shrimp paste. The pivot points in the graph represent the corresponding Taxonomy records in the NCBI database, the English name is spelled near the pivot point. The larger the abundance of a species is, the larger the circle of the fulcrum is. When a number of samples are plotted simultaneously, the relative abundance of different samples can be expressed in different colors by means of a small pie chart at the branches or nodes.

The proportion of *Proteobacteria* in the fermentation prophase was higher than in metaphase and anaphase. *Firmicutes* was most abundant in the metaphase stage, and present at the lowest levels in the prophase stage. *Bacteroidetes* showed a gradually decreasing trend across the three stages. The content of *Actinobacteria* and *Verrucomicrobia* differed little across the three stages. *Planctomycetes* and *Fusobacteria* were more abundant in the fermentation prophase than in metaphase or anaphase, and were present at extremely low levels in late fermentation (Table 2).

**TABLE 2:**
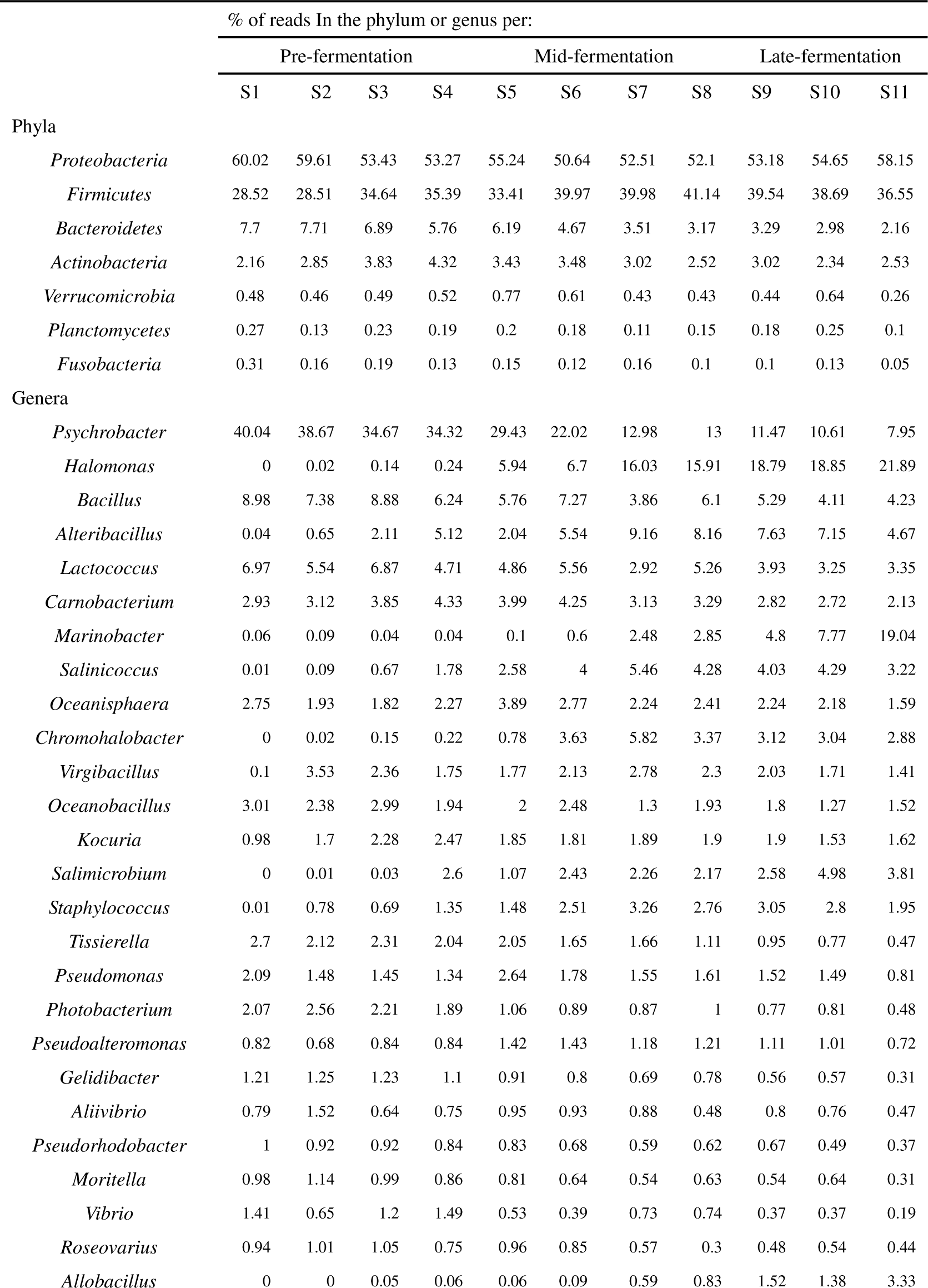
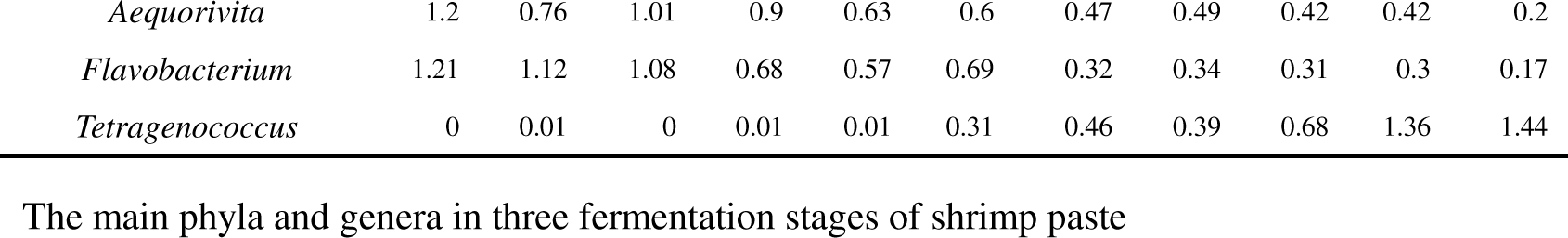
Summary of reads calculated from total phylum reads for variable assessed

At the genus level, *Psychrobacer* was the dominant microorganism, accounting for 23.78%, followed by *Halomonas* at 9.46% (Fig. 1). During the whole fermentation process, levels of *Psychrobacer* decreased (Fig.2), and the proportion of *Psychrobacer* was greater in fermentation prophase than in metaphase or anaphase (Table 2). *Halomonas* was not found in Stage 1 to Stage 4, but began to appear in Stage5, and gradually increased and stabilized (Fig. 2). This growth pattern may be related to the salt concentration of the shrimp paste during fermentation. Levels of *Bacillus* slowly increased from Stage 1 to Stage 3, began to decrease at Stage 4 and then remained essentially unchanged (Fig. 2). In general, *Bacillus* was most abundant in the pre-fermentation period (Table 2). *Alteribacillus* began to appear at a low level at Stage 2, gradually increased to Stage 8, then gradually reduced (Fig. 2). Its content was highest in fermentation metaphase than in the other two periods (Table 2). *Lactococcus* began to reduce after Stage 3 and then remained unchanged. After Stage 9 it decreased significantly (Fig. 2). The proportion of *Lactococcus* declined across the three fermentation phases (Table 2). *Carnobacterium* had the highest content mid-fermentation. *Marinobacter* levels were extremely low in the early stage of fermentation, and increased over time. *Salinicoccus, Chromohalobacter, Salimicrobium, Allobacillus*, and *Tetragenococcus* were almost nonexistent at the early stage and began to appear in the middle and late stage. There was little change in the content of *Oceanisphaera, Kocuria, Pseudomonas, Pseudoalteromonas*, and *Aliivibrio* in the three stages. The proportions of *Tissierella, Photobacterium, Gelidibacter, Pseudorhodobacterm, Moritella, Vibrio, Roseovarius, Aequorivita*, and *Flavobacterium* were higher in the early stage of fermentation than in the other stages. *Staphylococcus* content was highest in fermentation metaphase, followed by anaphase, and lowest in prophase.

**Figure 2:**
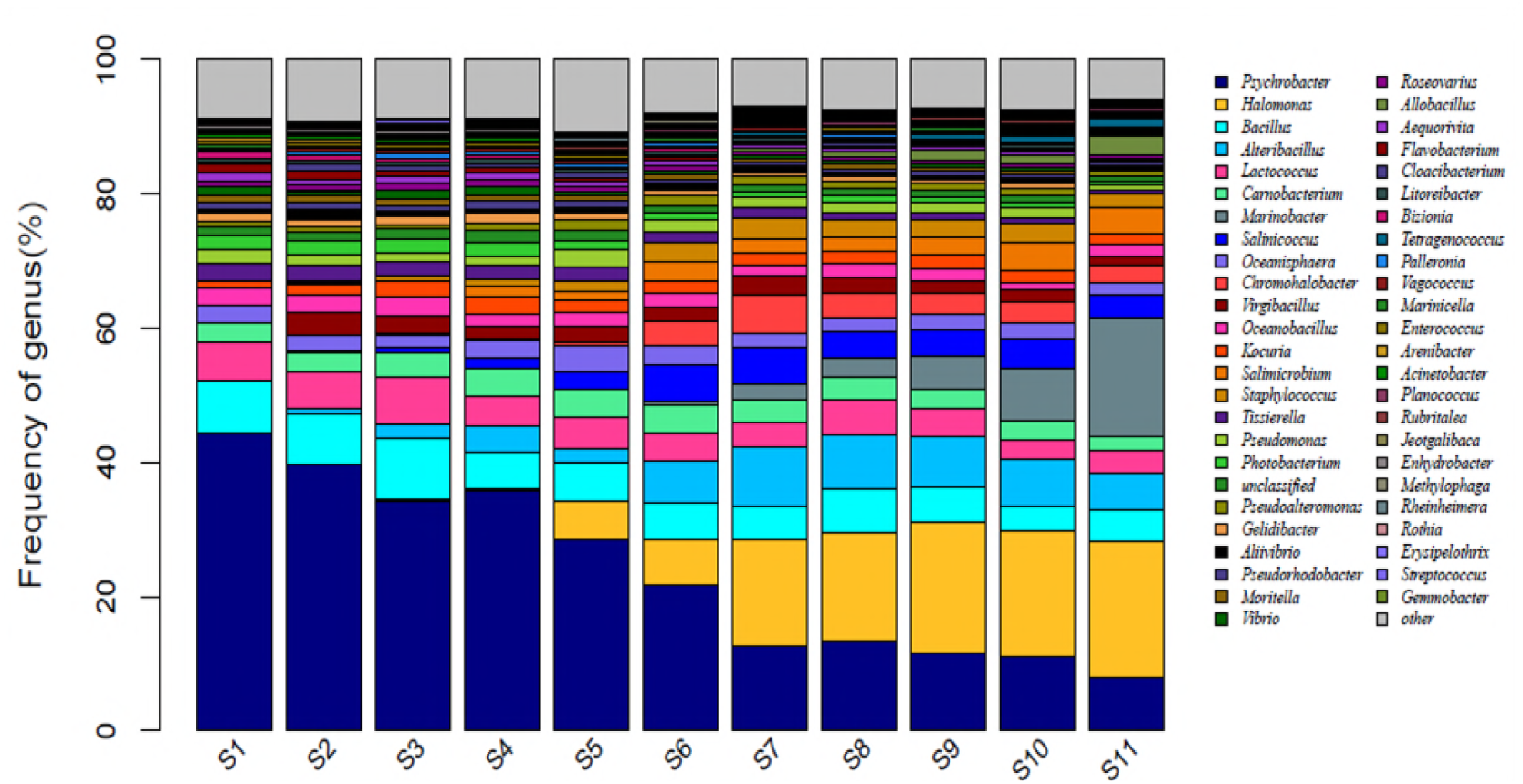
Assignment of shrimp paste at the genus level. The horizontal axis is the number of each sample, and the longitudinal axis is the relative abundance ratio. The color corresponds to the species name under the taxonomic level, and the width of different color blocks indicates the relative abundance ratio of differential species.

Principal coordinate analysis clustered the communities according to different stages (Fig. 3). Regardless of the community stage, there were no definitive splits in the microbiota of the shrimp paste. However, the most extreme outliers tended to be in Stage 11 in the fermentation anaphase. No statistical differences were found in operational taxonomic units at the genus level.

**Figure 3:**
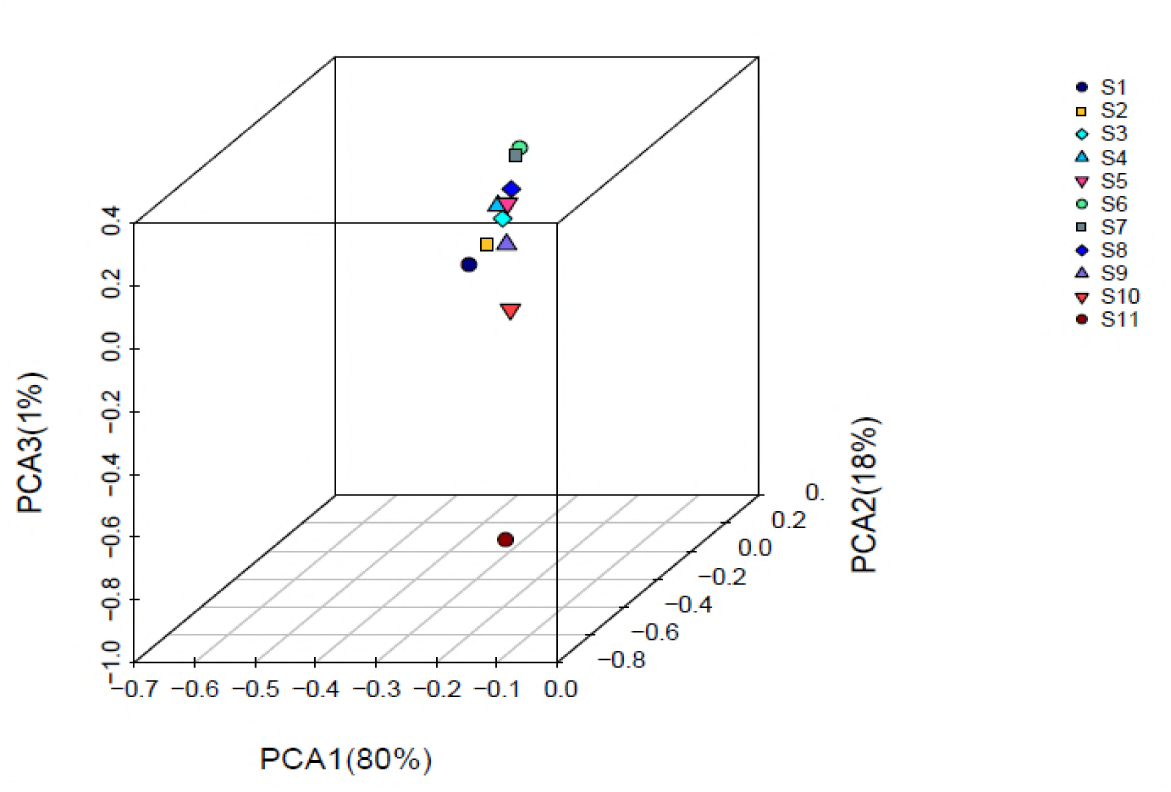
Principal coordinate analysis graphs of shrimp paste. Different colors represent different samples or different group samples in the graph, the higher the similarity between samples, the more likely to be aggregated in the graph.

## Discussion

In this study, high-throughput sequencing provided detailed insights into the complex microbiota of shrimp paste at different fermentation stages. We found that there were 30 phyla, 55 classes, 86 orders, 206 families, and 695 genera in the shrimp paste.

*Proteobacteria* was dominant, reaching 55.59%, and it was high at the pre-fermentation stage. *Proteobacteria* is a major group of gram-negative bacteria, which is widespread in humans (24), shrimp (25) and crabs (26–27). It is the dominant group in coastal areas and aquaculture ponds (28–32). The shrimp used in this study came from the Yellow Sea, where the major bacterial group is *Proteobacteria*, and therefore the dominance of *Proteobacteria* in the shrimp paste may be related to its source materials. The *Proteobacteria* is the largest group of bacteria, and it includes many pathogenic bacteria, such as *E. coli, Salmonella*, and *Helicobacter pylori* (33). *Proteobacteria* levels in food are mainly related to its freshness, and transport and storage hygiene conditions. Therefore, *Proteobacteria* may greatly influence the quality of shrimp paste, and it is necessary to strictly control their growth during its fermentation, production, and storage to standardize production.

*Psychrobacter* was the most abundant genus in the shrimp paste. It was present at highest levels in the early stage of fermentation. *Psychrobacter* is facultatively anaerobic and is common in high-salt foods (34). Once it appears in high-salt foods, it multiplies and can easily cause food spoilage, so it is key to the control of fermented foods (35–36). The gram-negative bacteria *Halomonas*, which also grows in high concentrations of salt (37–38), occurred mainly in late fermentation. Its presence has been suggested to be an indicator of hygiene problems in shrimp paste (39). *Psychrobacter* and *Halomonas* can produce lipolytic enzymes that promote the degradation of flavoring substances and play an important role in the formation of flavor (40–41). Across the three fermentation stages, *Bacillus* and *Lactococcus* generally presented a decreasing trend. *Bacillus* can produce butyric acid, which can promote the growth of less acid-tolerant spoilage microorganisms and thus contaminate the shrimp paste (42). *Lactococcus* can react with the alcohols, aldehydes, and ketones produced during fermentation to form a variety of new taste substances. In addition, various antibacterial substances such as organic acids, diacetyl, hydrogen peroxide, and bacteriocins produced by *Lactococcus* may prevent the growth of food spoilage bacteria and extend the shelf life of shrimp paste. *Lactococcus* may thus improve the functional attributes and shelf life of shrimp paste (43–45).

Several previous studies have focused on naturally fermented foods, including artisanal cheeses (46), Chinese liquor (47), and dairy products (48–49). In addition, there has been some research into the microbial diversity of shrimp culture environments, and the chemical composition and sensory attributes of shrimp paste [50]. However, there has not been a study of the microbial diversity of shrimp paste, which is a naturally fermented food. We addressed this research gap and found that the presence of bacteria corresponded mainly to four phyla (*Proteobacteria, Firmicutes, Actinobacteria*, and *Bacteroidetes*), and five major genera (*Psychrobacer, Halomonas, Bacillus, Alteribacillus*, and *Lactococcus*). These microbes all have a certain influence on the quality and flavor of shrimp paste.

We used high-throughput sequencing to investigate the microbiota of shrimp paste at different fermentation stages. Our analysis furthers understanding of the microbial community of shrimp paste and the relationship between microbial diversity and shrimp paste flavor. Our findings are of great significance to the technological control of shrimp paste. The standardization of production is important for ensuring the quality and safety of shrimp paste and improving its quality and nutritional value.

## Materials and Methods

### Sample collection and nucleic acid extraction

Shrimp paste samples were collected from Lianyungang, in Jiangsu Province. The shrimp paste was freshly produced in May 2017. During the production process, three stages of shrimp paste were analyzed. Samples were obtained during fermentation prophase, fermentation metaphase, and fermentation anaphase. The whole fermentation process lasted for 1 month and each stage took 10 days. The samples taken from fermentation prophase were labeled S1–S4, and the samples taken from fermentation metaphase and fermentation anaphase were defined as S5–S8 and S9–S11, respectively. DNA was extracted using an E.Z.N.A™ Mag-Bind Soil DNA Kit (OMEGA, USA).

### PCR amplification of the microbial community 16S rRNA genes

The DNA extracts were used as a template for PCR amplification according to the methods described for Qubit 2.0 (Life, USA). The V3-V4 region of bacterial 16S rRNA was amplified by PCR for high-throughput sequencing. The PCR reaction included two rounds of amplification, resulting in more specific and accurate results by sequencing at both ends. In the first round of amplification, the bacterial 16S rRNA gene V3-V4 region was amplified with the universal forward 341F (CCTACGGGNGGCWGCAG) and reverse 805R (GACTACHVGGGTATCTAATCC) primers. These primers contained a set of 6-nucleotide barcodes. The PCR mixture contained 15 μl 2×; Taq master mix, 1 μl bar-PCR primer F (10 uM), 1 μl primer R (10 uM), 10–20 ng genomic DNA, and ultra-pure H_2_O to give a final reaction volume of 30 μl. PCR amplification of the 16S rRNA V3-V4 regions was performed using a T100™ Thermal Cycler (BIO-RAD, USA). The amplification program was as follows: 1 cycle of denaturing at 94°C for 3 min, 5 cycles of denaturing at 94°C for 30 s, annealing at 45°C for 20 s, elongation at 65°C for 30 s, then 20 cycles of denaturing at 94°C for 20 s, annealing at 55°C for 20 s, elongation at 72°C for 30 s and a final extension at 72°C for 5 min. In the second round of amplification, the PCR mixture contained 15 μl 2×; Taq master mix, 1 μl primer F (10 uM), 1 μl primer R (10 uM), 20 ng genomic DNA, and ultra-pure H_2_O to give a final reaction volume of 30 μl. PCR amplification of the 16S rRNA V3-V4 regions was performed using a T100™ Thermal Cycler (BIO-RAD, USA). The amplification program was as follows: 1 cycle of denaturing at 95°C for 3 min, then 5 cycles of denaturing at 94°C for 20 s, annealing at 55°C for 20 s, elongation at 72°C for 30 s and a final extension at 72°C for 5 min. The PCR products were checked using electrophoresis in 1% (w/v) agarose gels in TBE buffer (Tris, boric acid, EDTA) stained with ethidium bromide and visualized under UV light. We used Agencourt AMPure XP beads (Beckman, USA) to purify the free primers and primer dimer species in the amplification product. Before sequencing, the DNA concentration of each PCR product was determined using a Qubit 2.0 kit and it was quality controlled using a bioanalyzer (Agilent, USA). The amplifications from each reaction mixture were pooled in equimolar ratios based on their concentrations.

### High-throughput sequencing and bioinformatics analysis

The V3-V4 region of bacterial 16S rRNA was sequenced on an Illumina MiSeq system (Illumina MiSeq, USA), according to the manufacturer’s instructions. Raw sequences were selected based on sequence length, quality, primer, and tag, and data were collected as follows. (i) The two short Illumina readings were assembled by PEAR (v.0.9.6) software according to the overlap and fastq files were processed to generate individual fasta and qual files, which could then be analyzed by standard methods. (ii) Sequences containing ambiguous bases and any longer than 480 bp were dislodged and those with a maximum homopolymer length of 6 bp were allowed. Sequences shorter than 200 bp were removed. (iii) All identical sequences were merged into one. (iv) Sequences were aligned according to a customized reference database. (v) The completeness of the index and the adaptor was checked and removed all of the index and the adaptor sequence. (vi) Noise was removed using the Pre.cluster tool. Chimeras were detected using Chimera UCHIME (v.4.2.40). We submitted the effective sequences of each sample to the RDP Classifier to identify bacterial and fungal sequences. Species richness and diversity statistics including coverage, chao1, ACE, Simpson and Shannon indexes were calculated using MOTHUR (v.1.30.1). A rarefaction curve was used to monitor results for sequencing abundance with the MOTHUR package. Principal coordinate analysis, measuring dissimilarities at phylogenetic distances based on weighted and unweighted Unifrac analysis, was performed using the QIIME suite of programs. According to the classification results, it was possible to determine the classification of the samples at the taxonomic level. The results indicated (i) which microorganisms were contained in the sample, and (ii) the relative abundance of each microorganism in the sample. The results at the species level can be analyzed by R(v3.2). We used MUSCLE (v.3.8.31) for multi-sequence alignment to obtain the alignment file, and FASTTREE (v.2.1.3) approximately-maximum-likelihood to construct the phylogenetic tree, inferring the order of the biological evolution of the sample. After analyzing the composition of the gene functions of the sequenced microbial genome, PICRUST (v.1.0.0) was used to analyze the differences in function between different samples and groups obtained by sequencing according to their functional gene composition.

## Acknowledgments

This work was supported by the Jiangsu Provincial Department of Science and Technology (BE2017316, BE2017326)

We thank Lianyungang Haiwa Food Co., Ltd. for the shrimp paste.

I wish to thank WANG Limei for guidance on the experimental and have made a great contribution to this article.

## References

[1] Jung W Y, Lee H J, Jeon C O. Halomonas garicola sp. nov., isolated from saeu-jeot, a Korean salted and fermented shrimp paste[J]. International journal of systematic and evolutionary microbiology, 2016, 66(2): 731–737. DOI 10.1099/ijsem.0.000784

[2] Duan S, Hu X, Li M, et al. Composition and Metabolic Activities of the Bacterial Community in Shrimp Paste at the Flavor-Forming Stage of Fermentation As Revealed by Metatranscriptome and 16S rRNA Gene Sequencings[J]. Journal of agricultural and food chemistry, 2016, 64(12):2591–2603. doi: 10.1021/acs.jafc.5b05826

[3] Mao X, Zhang J, Kan F, et al. Antioxidant production and chitin recovery from shrimp head fermentation with Streptococcus thermophilus[J]. Food Science and Biotechnology, 2013, 22(4):1023–1032. doi:10.1007/s10068-013-0179-5

[4] Mao X, Liu P, He S, et al. Antioxidant properties of bio-active substances from shrimp head fermented by Bacillus licheniformis OPL-007[J]. Applied biochemistry and biotechnology, 2013, 171(5):1240–1252. DOI :10.1007/s12010-013-0217-z

[5] Pongsetjul J, Benjakul S, Sampavapol P, et al. Chemical compositions, sensory and antioxidative properties of salted shrimp paste (Ka-pi) in Thailand[J]. International Food Research Journal, 2015, 22(4):1454–1465

[6] Halder S K, Adak A, Maity C, et al. Exploitation of fermented shrimp-shells hydrolysate as functional food: assessment of antioxidant, hypocholesterolemic and prebiotic activities[J]. Indian Journal of Experimental Biology, 2013, 51(12):924–934

[7] Faithong N, Benjakul S. Changes in antioxidant activities and physicochemical properties of Kapi, a fermented shrimp paste, during fermentation[J]. Journal of Food Science & Technology, 2014, 51(10):2463–2471. DOI 10.1007/s13197-012-0762-4

[8] Pongsetkul J, Benjakul S, Sampavapol P, et al. Chemical compositions, sensory and antioxidative properties of salted shrimp paste (Ka-pi) in Thailand.[J]. International Food Research Journal, 2015.

[9] Sun G Y, Zuo Y P. Fermentation Technology and Research Progress of Shrimp Paste[J]. China Condiment, 2013.

[10] Soon W W, Hariharan M, Snyder M P. High-throughput sequencing for biology and medicine[J]. Molecular systems biology, 2013, 9(1):640–648. doi:10.1038/msb.2012.61

[11] Alessandria V, Ferrocino I, De Filippis F, et al. Microbiota of an Italian Grana-Like cheese during manufacture and ripening, unraveled by 16S rRNA-Based approaches[J]. Applied and environmental microbiology, 2016, 82(13): 3988–3995. DOI:10.1128/AEM.00999-16.

[12] Wey J K, Jürgens K, Weitere M. Seasonal and successional influences on bacterial community composition exceed that of protozoan grazing in river biofilms[J]. Applied and environmental microbiology, 2012, 78(6): 2013–2024. doi:10.1128/AEM.06517-11

[13] Robertson C E, Baumgartner L K, Harris J K, et al. Culture-independent analysis of aerosol microbiology in a metropolitan subway system[J]. Applied and environmental microbiology, 2013, 79(11): 3485–3493. DOI:10.1128/AEM.00331-13

[14] Ercolini D. High-throughput sequencing and metagenomics: moving forward in the culture-independent analysis of food microbial ecology[J]. Applied and environmental microbiology, 2013, 79(10):3148–3155. doi:10.1128/AEM.00256-13

[15] Li X R, Ma E B, Yan L Z, et al. Bacterial and fungal diversity in the starter production process of Fen liquor, a traditional Chinese liquor[J]. Journal of Microbiology, 2013, 51(4):430–438. DOI 10.1007/s12275-013-2640-9

[16] Wolfe B E, Button J E, Santarelli M, et al. Cheese rind communities provide tractable systems for in situ and in vitro studies of microbial diversity[J]. Cell, 2014, 158(2):422–433. http://dx.DOI.o/10.1016/j.cell.2014.05.041

[17] Gao J, Gu F, He J, et al. Metagenome analysis of bacterial diversity in Tibetan kefir grains[J]. European Food Research and Technology, 2013, 236(3):549–556. DOI 10.1007/s00217-013-1912-2

[18] Di Bella J M, Bao Y, Gloor G B, et al. High throughput sequencing methods and analysis for microbiome research[J]. Journal of microbiological methods, 2013, 95(3):401–414

[19] Jung J Y, Lee S H, Lee H J, et al. Microbial succession and metabolite changes during fermentation of saeu-jeot: traditional Korean salted seafood[J]. Food microbiology, 2013, 34(2): 360–368.

[20] Rezaee S, Farahmand H, Nematollahi M A. Genetic diversity status of Pacific white shrimp (Litopenaeus vannamei) using SSR markers in Iran[J]. Aquaculture International, 2016, 24(2):479–489.DOI 10.1007/s10499-015-9939-y

[21] Vaseeharan B, Rajakamaran P, Jayaseelan D, et al. Molecular markers and their application in genetic diversity of penaeid shrimp[J]. Aquaculture international, 2013, 21(2):219–241. DOI10.1007/s10499-012-9582-9

[22] Rajakumaran P, Vaseeharan B. Survey on Penaeidae shrimp diversity and exploitation in south east coast of India[J]. Fisheries and Aquaculture Journal, 2015. http://dx.DOI.org/10.4172/2150-3508.1000103

[23] Jinap S, Ilyanur A R, Tang S C, et al. Sensory attributes of dishes containing shrimp paste with different concentrations of glutamate and 5′-nucleotides.[J]. Appetite, 2010, 55(2):238–244.

[24] Qin N, Yang F, Li A, et al. Alterations of the human gut microbiome in liver cirrhosis[J]. Nature, 2014, 513(7516):59–78. doi:10.1038/nature13568

[25] Xiong J, Wang K, Wu J, et al. Changes in intestinal bacterial communities are closely associated with shrimp disease severity[J]. Applied microbiology and biotechnology, 2015, 99(16):6911–6919.DOI 10.1007/s00253-015-6632-z

[26] Li S, Sun L, Wu H, et al. The intestinal microbial diversity in mud crab (Scylla paramamosain) as determined by PCR-DGGE and clone library analysis[J]. Journal of applied microbiology, 2012, 113(6):1341–1351. doi:10.1111/jam.12008

[27] Zhang M, Sun Y, Chen L, et al. Symbiotic bacteria in gills and guts of Chinese mitten crab (Eriocheir sinensis) differ from the free-living bacteria in water[J]. PloS one, 2016, 11(1):e0148135–e0148147. doi:10.1371/journal.pone.0148135

[28] Gilbert J A, Steele J A, Caporaso J G, et al. Defining seasonal marine microbial community dynamics[J]. The ISME journal, 2012, 6(2):298–308.doi:10.1038/ismej.2011.107

[29] Zhao, Yanting, et al. “Metagenomic analysis revealed the prevalence of antibiotic resistance genes in the gut and living environment of freshwater shrimp.” Journal of hazardous materials 350 (2018): 10–18.

[30] Fan, Limin, et al. “Methanogenic community compositions in surface sediment of freshwater aquaculture ponds and the influencing factors.” Antonie van Leeuwenhoek 111.1 (2018): 115–124. https://DOI.org/10.1007/s10482-017-0932-5

[31] Cornejo-Granados, Fernanda, et al. “Microbiome of Pacific Whiteleg shrimp reveals differential bacterial community composition between Wild, Aquacultured and AHPND/EMS outbreak conditions.” Scientific reports 7.1 (2017): 11783. doi:10.1038/s41598-017-11805-w

[32] Zhang, Qing, et al. “Temporal heterogeneity of prokaryotic micro-organism communities in sediment of traditional freshwater cultured fish ponds in Southwest China.” Biotechnology & Biotechnological Equipment 32.1 (2018): 102–108. https://DOI.org/10.1080/13102818.2017.1400403

[33] Kilinc B, Besler A. Seafood toxins and poisonings[J]. Turkish Journal of Aquatic Sciences, 2015, 30(1): 35–52. doi: 10.18864/iujfas.60084

[34] Wu G, Zhang X, Wei L, et al. A cold-adapted, solvent and salt tolerant esterase from marine bacterium Psychrobacter pacificensis[J]. International journal of biological macromolecules, 2015, 81: 180–187.

[35] Stellato G, De Filippis F, La Storia A, et al. Coexistence of lactic acid bacteria and potential spoilage microbiota in a dairy processing environment[J]. Applied and environmental microbiology, 2015, 81(22): 7893–7904.doi:10.1128/AEM.02294-15.

[36] Ferrocino I, Bellio A, Romano A, et al. RNA-based amplicon sequencing reveals microbiota development during ripening of artisanal versus industrial Lard d’Arnad[J]. Applied and environmental microbiology, 2017, 83(16): e00983–17. https://DOI.org/10.1128/AEM.00983-17.

[37] Burch A Y, Finkel O M, Cho J K, et al. Diverse microhabitats experienced by Halomonas variabilis on salt-secreting leaves[J]. Applied and environmental microbiology, 2013, 79(3): 845–852. doi:10.1128/AEM.02791-12

[38] Kaye J Z, Baross J A. Synchronous effects of temperature, hydrostatic pressure, and salinity on growth, phospholipid profiles, and protein patterns of four Halomonas species isolated from deep-sea hydrothermal-vent and sea surface environments[J]. Applied and environmental microbiology, 2004, 70(10): 6220–6229. doi: 10.1128/AEM.70.10.6220–6229.2004

[39] Maoz A, Mayr R, Scherer S. Temporal stability and biodiversity of two complex antilisterial cheese-ripening microbial consortia[J]. Applied and Environmental Microbiology, 2003, 69(7): 4012–4018. doi: 10.1128/AEM.69.7.4012–4018.2003

[40] Kim S, Wi A R, Park H J, et al. Enhancing extracellular lipolytic enzyme production in an arctic bacterium, Psychrobacter sp. ArcL13, by using statistical optimization and fed-batch fermentation[J]. Preparative Biochemistry and Biotechnology, 2015, 45(4): 348–364. doi: 10.1080/10826068.2014.940964

[41] Hinrichsen L L, Montel M C, Talon R. Proteolytic and lipolytic activities of Micrococcus roseus (65), Halomonas elongata (16) and Vibrio sp.(168) isolated from Danish bacon curing brines[J]. International journal of food microbiology, 1994, 22(2-3): 115–126.

[42] Duniere L, Xu S, Long J, et al. Bacterial and fungal core microbiomes associated with small grain silages during ensiling and aerobic spoilage[J]. BMC microbiology, 2017, 17(1): 50. DOI 10.1186/s12866-017-0947-0

[43] Peng K, Jin L, Niu Y D, et al. Condensed tannins affect bacterial and fungal microbiomes and mycotoxin production during ensiling and upon aerobic exposure[J]. Applied and environmental microbiology, 2018, 84(5): e02274–17. https://DOI.org/10.1128/AEM.02274-17.

[44] Minervini F, Conte A, Del Nobile M A, et al. Dietary fibers and protective lactobacilli drive burrata cheese microbiome[J]. Applied and environmental microbiology, 2017, 83(21): e01494–17. https://DOI.org/10.1128/AEM.01494-17.

[45] Frantzen C A, Kleppen H P, Holo H. Lactococcus lactis Diversity in Undefined Mixed Dairy Starter Cultures as Revealed by Comparative Genome Analyses and Targeted Amplicon Sequencing of epsD[J]. Applied and environmental microbiology, 2018, 84(3): e02199–17. https://DOI.org/10.1128/AEM.02199-17.

[46] Quigley L, O’Sullivan O, Beresford T P, et al. High-throughput sequencing for detection of subpopulations of bacteria not previously associated with artisanal cheeses[J]. Applied and environmental microbiology, 2012, 78(16): 5717–5723. doi:10.1128/AEM.00918-12

[47] Wang X, Du H, Zhang Y, et al. Environmental Microbiota Drives Microbial Succession and Metabolic Profiles during Chinese Liquor Fermentation[J]. Applied and environmental microbiology, 2018, 84(4): e02369–17. https://DOI.org/101128/AEM.02369-17.

[48] Cruciata M, Gaglio R, Scatassa M L, et al. Formation and characterization of early bacterial biofilms on different wood typologies applied in dairy production[J]. Applied and environmental microbiology, 2018, 84(4): e02107–17. https://DOI.org/10.1128/AEM.02107-17.

[49] Muhammed M K, Kot W, Neve H, et al. Metagenomic analysis of dairy bacteriophages: Extraction method and pilot study on whey samples derived from using undefined and defined mesophilic starter cultures[J]. Applied and environmental microbiology, 2017, 83(19): e00888–17. https://doi.org/10.1128/AEM00888-17.

[50] Pilapil A R, Neyrinck E, Deloof D, et al. Chemical quality assessment of traditional salt-fermented shrimp paste from Northern Mindanao, Philippines[J] Journal of the Science of Food & Agriculture, 2015, 96(3):933–938. DOI 10.1002/jsfa.7167.

